# Comparison of gene expression patterns of *Kappaphycus alvarezii* (Rhodophyta, Solieriaceae) under different light wavelengths and CO_2_ enrichment

**DOI:** 10.1101/188250

**Authors:** Thien Vun Yee, Kenneth Francis Rodrigues, Clemente Michael Wong Vui Ling, Wilson Yong Thau Lym

## Abstract

Transcriptomes associated with the process of photosynthesis and carbon fixation have offered insights into the mechanism of gene regulation in terrestrial plants, however limited information is available as far as macroalgae are concerned. Intertidal red alga, *Kappaphycus alvarezii* is exposed to different wavelengths of light in their lives as light quantity and quality changes at different depths in seawater. This investigation aims to study the underlying mechanisms associated with photosynthesis and carbon fixation under specific light qualities and CO_2_ enrichment. Light regulation of gene expression has not been previously described for red algae. By using next generation sequencing, transcriptome profiling of *K. alvarezii* generated 76,871 qualified transcripts with a mean length of 979bp and a N50 length of 1,707bp and 55.83% transcripts were annotated on the basis of function. Blue, green and red light all have demonstrated roles in modulating light responses, such as changes in gene expression. Here we analysed the effects of light regulation on four selected photosynthesis aspects (light-harvesting complex, phycobilisomes, photosystems and photoreceptors). We observed that light-regulated gene expression in this species is not a single light response and different light qualities are transduced to regulate the same metabolic pattern. The carbon fixation pathway was analysed and key genes encoding enzymes involved in the carbon fixation pathway such as ppc, pepc, prk, pgk, ppdk, provided that unequivocal molecular evidence that most of the C_3_ and C_4_ pathway genes were actively transcribed in *K. alvarezii*. In addition to this the CO_2_ induced transcriptome suggested the possibility of shifting carbon metabolism pathway after acclimation to increased level of CO_2_. Impact of CO_2_ enrichment on the cultures has provided new insight into the response to rising CO_2_.

## Introduction

Transcriptome shifts associated by the light signals perceived by photoreceptors have offered insights into the gene regulation in higher plants, however limited information is available as far as macroalgae under light treatment are concerned. Red alga, *Kappaphycus alvarezii* grows in intertidal zone where they may face irradiance environments depleted in both light quantity and quality as the light spectral distribution changes at different depths in seawater. Intertidal seaweeds exposed to light spectrum which is similar as in air during low tide and blue light becomes predominant where the seaweeds grow under seawater of 2-5 m in depth at high tide (Dring, 1981). Seaweeds can acclimate to the changes of light quantity and qualities by employing efficient light-harvesting mechanisms, such as increase the quantity of photosynthetic pigments, change the ratio of accessory pigments to chlorophyll *a* (Lobban and Harrison, 1994). Red alga, *Chondrus crispus* have been reported to have increased phycoerythrin content and the efficiencies of photosynthesis increased with depth (Sagert *et al*., 1997).

Recently, insights in the light-mediated physiological responses and molecular mechanisms have been gained in brown alga, *Saccharina japonica* (Deng *et al*., 2012; Wang *et al*., 2013). Expression profiling of their researches indicate that light induces profound gene expression changes in *S. japonica*. These light responsive genes include many transcription factors and fall into various functional categories mainly involved in photomorphogenesis, circadian clock function, photoreactivation, photosynthetic carbon, metabolism and biosynthesis. Wang *et al*. (2013) suggested that promotion of metabolism and growth in kelps under blue light predominating environment was attributed to both reduction of red light and increase of blue light illumination. Thus, it suggested that different light qualities may play significant roles in the lives of red algae.

The transcriptome of *K. alvarezii* was first sequenced by Wu *et al*. (2014) together with 19 Phaeophyceae (brown algae) and other 20 Rhodophyceae (red algae), provide a broad range of algal transcriptome information that could contribute to algal genetic and biological study. Then, Song *et al*. (2014) performed de novo transcriptome sequencing of *K. alvarezii* to investigate the mechanisms underlying the biosynthesis of carrageenan in Family Solieriaceae (*Betaphycus gelatinus*, *K. alvarezii* and *Eucheuma denticulatum*). They have identified 861 KEGG orthologs which might contain the main genes regulating the biosynthesis of carrageenan with different types and possessions. More recently, the transcriptome of *K. alvarezii* was profiled by Zhang *et al*. (2015). The results elucidate some genetic information and reveal many important metabolic pathways in *K. alvarezii*. For instance, the significantly enriched pathways include selenocompound metabolism (ko00450) and sulphur metabolism (ko00920) which were probably related to the alga’s resistance to grazers and expelling of toxins (Lobanov *et al*., 2007).

No clear evidence for C_3_ or C_4_ photosynthetic pathways have been found in the *K. alvarezii*. The occurrence photosynthetic pathways in *K. alvarezii* are therefore still in question and await further research. Many studies reported that the operation of C_3_ pathway is predominant in algae (Beer *et al*., 1986; Tsuji *et al*., 2009); however, it has been demonstrated that C_4_ photosynthesis is found in some eukaryotic algae (Derelle *et al*., 2006; Leliaert *et al.,* 2012). The study of *Ulva prolifera* and *Chara contraria* on their primary photosynthetic carbon metabolism revealed that some algae possess both C_3_ and C_4_ pathways and may alter their carbon metabolism pursuant to the environment (Keeley, 1999; Xu *et al*., 2012). The alterations of photosynthetic pathways under environmental changes probably contributing to the adaptation of plants to environmental stress (Ehleringer *et al*., 1997). A submerged aquatic plant, *Hydrilla verticallata*, was reported to change its photosynthetic pathways from C_3_ to C_4_ under conditions of CO_2_ deficiency (Reiskind *et al*., 1997).

In this study, we used next generation sequencing technology to profile the transcriptome of red alga, *K. alvarezii*, with the aim to characterise its functional genome under specific light qualities and CO_2_ enrichment. This investigation will enhance our understanding of molecular mechanisms underlying light-induced responses in lower plants as well as facilitate our understanding in inorganic carbon fixation in red algae. In addition, this study would improve our understanding of the impact of changing light conditions on *K. alvarezii* and the potential mechanism of light adaptation of this red alga.

### Materials and methods

#### Seaweed materials and culture conditions

Seedlings of *Kappaphycus alvarezii* (var. *tambalang* ‘giant’) were obtained from Biotechnology Research Institute (BRI), Universiti Malaysia Sabah. The seaweed was originally collected from Pulau Sebangkat (4°33′ 31″N, 118°39′49″E), Semporna, Sabah. The young seedlings were cultured under laboratory conditions (Yong *et al*. 2014). The experimental samples were selected from healthy and disease-free explants. Three replicates consisted of five algal tips were cut to a total length of 2 cm and used for each experimental condition. The algal tips were cultured in the Fernbach culture vessel with 800 mL artificial seawater (Fluval marine salt, 36 g L^-1^, salinity 31.4 ppt) enriched with 50 % Provasoli’s enriched seawater (PES) media. For light treatment, the cultures grown under 75 μmol photons m^−2^ s^−1^ white light (WL), blue light (BL) (wavelength = 492-455 nm), green light (GL) (wavelength = 577-492 nm) and red light (RL) (wavelength = 780-622 nm), respectively. Cultures were carried out at 25±1.0°C with 18 h light and 6 h dark cycle. Light-emitting diodes (LEDs) were used as light sources. Detected irradiances were measured with a digital light meter (Kyoritsu, Japan). For CO_2_ treatment, the cultures were illuminated by white LED, providing an irradiance of 75 μmol photons m^−2^ s^−1^, on a 18 h light and 6 h dark cycle. CO_2_ was provided 500 mL per day and controlled by supplying chambers with air/CO_2_ mix. Provision of 500 mL of CO_2_ per day was proved to achieve the highest growth rate of *K. alvarezii* (Barat, 2011). To avoid overly acidifying cultures, supplemental CO_2_ was only supplied during photoperiod. Temperature was maintained at 25±1.0°C. The cultures from both treatments were collected for RNA extraction after 14 days experimental period.

#### RNA isolation and preparation of cDNA library

Total RNA was extracted using QIAGEN RNeasy Plant Mini Kit (QIAGEN, Germany) according to manufacturer’s protocol. The experimental samples were wash with DEPC-treated water and then quickly frozen with liquid nitrogen and ground to a fine powder using RNase-free, chilled mortar and pestle. Typically, RNA samples with an A260/A280 ratio between 1.8 – 2.0 and A260/A230 ratio between 1.8 – 2.1 and RIN number of 8 and above were recommended to be sufficiently pure for further library construction.

Five cDNA libraries were generated with messenger RNA (mRNA) isolation and cDNA synthesis from five RNA samples were performed using NEBNext Ultra RNA Library Prep Kit for Illumina according to manufacturer’s protocol. In brief, the mRNA was purified from total RNA using poly-T oligo-attached magnetic beads and sheared into short segments of about 200 bp. Using the cleaved short RNA fragments as templates, first strand cDNAs were synthesised by random hexamer-primers and reverse transcriptase. The second strand cDNA was synthesised using DNA polymerase I and RNase H. The purified double strand cDNAs were subjected to end repair and NEBNext adaptor ligation. The resulting libraries were then loaded onto a single lane of the flow cell, and 209 cycles on the Illumina HiSeq 2000 platform (Illumina, USA) performed by Malaysian Genomics Resource Centre (MGRC). The high quality reads were deposited at the NCBI Short Read Archive (SRA) with the accession number: SRR2757332 (Green), SRR2757333 (Blue), SRR2757334 (CO_2_), SRR2757335 (White) and SRR2757337 (Red).

#### Quality control and de novo transcriptome assembly

Sequencing reads from the Illumina sequencer were exported in FASTQ format with the corresponding Phred quality scores. First, the quality of the sequencing raw reads was evaluated with FastQC v0.11.2 (www.bioinformatics.babraham.ac.uk/projects/fastqc). Strict reads filtering was performed before assembly: (i) removing sequencing adaptor sequences; (ii) filtering the reads containing unknown nucleotides (Ns); (iii) removing the reads with more than 10% bases below Q20 sequencing quality.

The obtained clean reads were subjected to transcriptome de novo assembly using Trinity (Grabherr *et al*., 2011). Briefly, Inchworm first assembles reads by searching for paths in a 25 bp K-mer graph, resulting in a collection of linear contigs. Next, Chrysalis clusters the reads with certain length of overlap and constructs a De Bruijn graph for each cluster. Finally Butterfly reconstructs plausible full-length, linear transcripts by reconciling the individual de Bruijn graphs generated by Chrysalis. After generating final assembly, TGIR Gene Indices clustering tools (TGICL) software (Pertea *et al*., 2003) was used to remove redundant sequences.

#### Transcriptome analysis

The assembled sequences were blastx searched against the UniProt protein databases (The UniProt Consortium, 2015) and Kyoto Encyclopaedia of Genes and Genomes (KEGG) database (Kanehisa *et al*., 2008) with 10^-5^ *E*-value cutoff. The sequences were then functionally clustered using Blast2GO (Conesa *et al.,* 2005) to get GO annotation and further classification by WEGO (Ye *et al*., 2006) for all assembled sequences.

#### Identification of differentially expressed genes (DEGs)

RSEM was used to estimate transcripts abundance (Li and Dewey, 2011) and differential expression between two groups was determined by software edgeR (Robinson *et al*., 2010). In order to analysing differential expression, the transcripts with adjusted *p*-value<1e-3 and log_2_ fold change of two were extracted and clustered according to their patterns of differential expression across the sample. All the DEGs were mapped to the terms in KEGG database and searched for significantly enriched KEGG pathways compared to the whole transcriptome background.

#### Quantitative real-time PCR analysis

Total RNA was extracted with QIAGEN RNeasy Plant Mini Kit (QIAGEN, Germany). The cDNA synthesis was carried out with QuantiTect Reverse Transcription kit (QIAGEN, Germany). The 18S rDNA gene (U25437) was chosen as internal control for normalisation of real-time PCR data. Primers for amplification of targeted genes (Supplementary Table S1) and 18S rDNA gene were designed using Primer3 software (http://frodo.wi.mit.edu/primer3) aiming for an amplicon length of 80 – 200 bp, a GC content of 40-60%, a primer length of 18 – 30 bp and a primer melting temperature of 60 – 65°C. Primers were assessed for melting temperature, hairpins and primer dimers using the web-based tool Beacon Designer (http://free.premierbiosoft.com). The real-time quantitative PCR was performed with QuantiNova SYBR green master mix (QIAGEN, Germany) on the Eco Real-Time PCR System (Illumina). Real-time PCR was performed in volume of 20 μL, and cycling conditions were 95°C for 2 min; 40 cycles of 95°C for 5 s and 60°C for 10 s. The PCR reactions were repeated in triplicates together with negative control and data were analysed using the 2^-ΔΔCT^ method.

## Results

### Transcriptome sequencing and de novo assembly

The sequencing run generated a total of 31.89 Gb raw paired-end reads. After trimming adapters and filtering out low quality reads, over 25 Gb clean reads (∼79%) were retained for assembly and further analysis. We generated 76,871 assembled transcripts by Trinity, which have lengths ranging from 300 to 17,257 bp (Fig. 1) and an N50 of 1,707 bp (Table 1). There were no ambiguous bases within the assembled sequences. A large number of the reads (93.51%) aligned back to the transcripts and average read depth of 256.47. The reads that did not map back to the assembled transcripts corresponded to either shorter than 300 bp or failed to match with any clean read.

**Figure 1:**
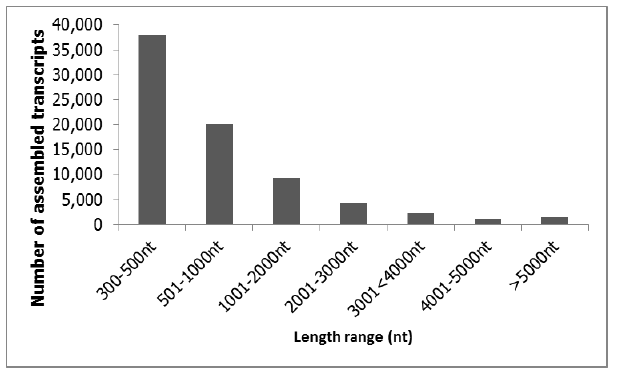
Graphical length distribution summary of transcripts identified in transcriptome data.

**Table 1:** Assembly results of *K. alvarezii* transcriptome generated from Trinity.

|  | Trinity |
| --- | --- |
| Total transcripts | 76871 |
| total length (bp) | 75,239,915 |
| Mean length (bp) | 979 |
| N50 length (bp) | 1,707 |
| The shortest transcripts (bp) | 300 |
| The longest transcripts (bp) | 17,257 |
| Transcripts $\geq 500$ bp (%) | 50.63% |
| Transcripts $\geq 1000$ bp (%) | 24.44% |
| GC (%) | 51.17 |
| Annotation rate (%) | 55.83 |

## Functional annotation of *K. alvarezii* transcriptome

Blastx was used to map assembled transcripts against the protein databases UniProt, KEGG and COG with a cutoff *E*-value of 10^-5^. The top hit from each of these assembled transcripts comparisons was used as the annotation reference for the respective transcripts. To sum up, 42,915 transcripts were assigned to putative functions, accounting 55.83% of the total assembled sequences (Supplementary Table S2). With UniProt annotation, 23,701 sequences had perfect matches with *E*-value <10^-45^ and 21.42% of the matches had similarity over 95% (Fig. 2a, b). The species distribution showed that only 13.45% had top matches to *Chondrus crispus* and next top matching species was *Phaeodactylum tricornutum*, accounting for 3.61% (Fig. 2c).

**Figure 2:**
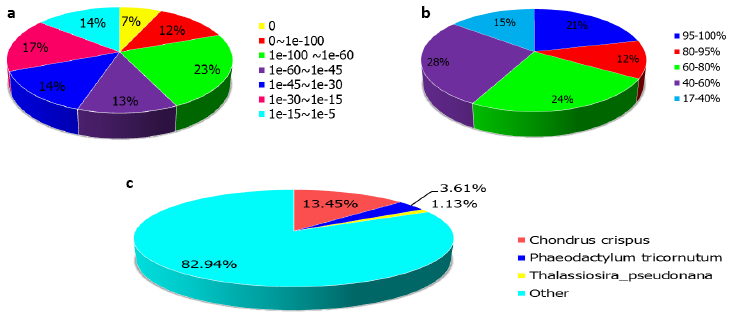
Characteristics of similarity search of the assembled sequences against the Uniprot databases. **a** E value distribution of blast hits for each unique sequence with E value ≤ 10-5. **b** Similarity distribution of the top blast hits for each sequence. **c** Species distribution of the total homologous sequences with E value ≤ 10-5. The first hit of each sequence was used for statistics.

There were 45 transcripts encoding light harvesting proteins and 72 gene tags belonging to photosynthetic pigments including chlorophyll, carotenoids (beta‐ and zeta-carotene), phycobilisome, phycocyanin and phycoerythrin. Three are types photoreceptors were identified including one cryptochrome, two blue-light receptor (PHR2) and four phytochrome protein. One phytochrome protein had significant homology to the phytochrome-like protein 2, a photoreceptor which exists in two forms that are reversibly interconvertible by light (Kaneko *et al*., 1995). In addition, a total of 38 gene tags belonging to photosystem proteins (I and II). The main subunits of PS I, PsaA, PsaB and PsaC, PS I chlorophyll A apoprotein, the Ycf3 and Ycf4 protein domain are identified. Out of 38 photosystem proteins, 23 of them are photosystem II proteins including D1, D2, CP43, CP47, PsbP, PS II reaction center protein M). There were 307 gene tags belonging to protein kinase (PK) family, including serine/threonine PK, calcium-dependent PK and receptor-like PK. A total of 59.93% PKs in *K. alvarezii* were highly homologous to those in *Chondrus crispus*.

To facilitate the organization of the *K. alvarezii* transcripts into putative functional groups, Gene Ontology (GO) terms were assigned using WEGO software. A total of 28,079 annotated transcripts were further categorized to the three main GO domains: biological processes, cellular components and molecular function, including 20,847 sequences at term “biological process”, 14,291 sequences at term “cellular component” and 41,698 sequences at term “molecular function” (Supplementary Fig. S3).

Functional and pathway analysis of the transcripts of *K. alvarezii* were carried out using the KEGG pathway database (Kanehisa *et al*., 2004). A total of 10,460 transcripts had assigned to 278 KEGG pathways (Table 2). Category “metabolic pathways” had the largest numbers of transcripts among all the categories, followed by the categories of “biosynthesis of secondary metabolites”, “biosynthesis of amino acids” and “carbon metabolism”.

**Table 2.**
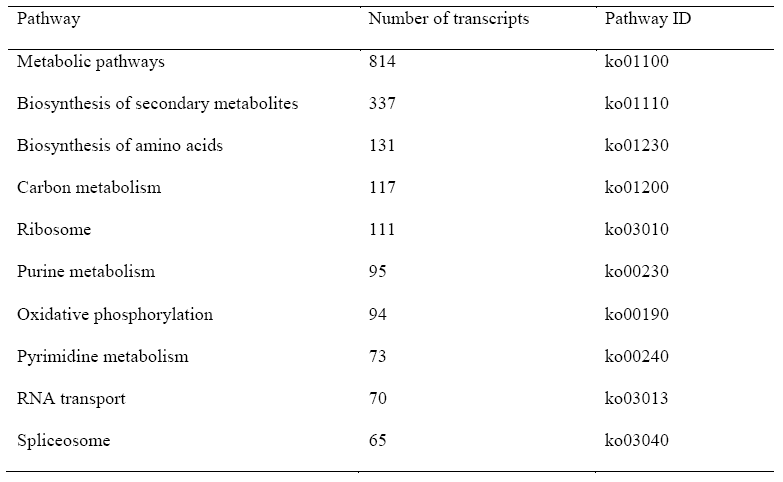
The top 10 pathways identified in the *Kappaphycus* transcriptome. Pathway

Both C3 and C4 photosynthesis genes were found in *K. alvarezii* by transcriptome sequencing. In total, 97 transcripts assigned to all the essential enzymes in both C3 and C4 carbon fixations were identified (Supplementary Table S4). Some key genes of enzymes involved in the carbon fixation pathway in *K. alvarezii* were discovered, such as aspartate aminotransferase (ast), phosphoribulokinase (prk), pyruvate orthophosphate dikinase (ppdk), phosphoglycerate kinase (pgk), malate dehydrogenase (mdh), phosphoenolpyruvate carboxylase (ppc) and phosphoenolpyruvate carboxykinase (pck), which provided unequivocal molecular evidence that most of the C_3_ and C_4_ pathway genes were actively transcribed in *K. alvarezii*.

### Identification of DEGs

The high quality reads from individual samples were mapped to the transcriptome database with the mapping percentage ranged from 84 to 86% (Table 3). The lower mapping rate reflects the low transcriptome sequencing depth and the high heterozygosity found in *K. alvarezii.*

**Table 3.** Paired-end read mapping statistics of transcripts identified in transcriptome data. Sample WL BL GL RL C02E* Total reads pair 30,115,700 22,796,939 27,106,635 23,380,754 24,122,534 Total mapped

| Sample | WL | BL | GL | RL | C02E* |
| --- | --- | --- | --- | --- | --- |
| Total reads pair | 30,115,700 | 22,796,939 | 27,106,635 | 23,380,754 | 24,122,534 |
| Total mapped<br>reads pair | 25,435,206 | 19,493,673 | 23,083,936 | 19,978,487 | 20,784,761 |
| Percentage of<br>mapped reads<br>pair (%) | 84.46 | 85.51 | 85.16 | 85.45 | 86.16 |

\*CO2E indicated CO<sub>2</sub> enriched cultures.

The read normalisation was performed using TMM methods (Robinson and Oshlack, 2011) which implemented in the edgeR Bioconductor package. Setting FDR ≤10^-3^ and log2 fold change of two as the cutoff, 11,769 DEGs were obtained (Supplementary Table S5). The results were validated by qPCR (data not shown), confirming the accuracy of RNA-Seq in mRNA quantification. A total of 7,962 DEGs (67.65%) were successfully annotated through the sequence similarity searching against the UniProt databases with a significance of E-value ≤ 10^-5^.

The most DEGs were found to be down-regulated between every two samples. For the light treatment, the most DEGs were detected between GL and WL, with 390 DEGs upregulated and 908 DEGs downregulated (Supplementary Fig. S5). If mRNA abundance of a gene was similar between BL, GL and WL while higher than RL, the DEG was interpreted to be RL-downregulated. If the expression level of a gene was higher or lower under WL than under both BL, GL and RL, the DEG was suggested to be either BL‐, GL‐ or RL-regulated. Based on the gene expression patterns among the four lights (Table 4), a total of 5,967 DEGs were designated to four categories: BL-regulated, GL-regulated, RL-regulated and either BL‐ or GL‐ or RL-regulated (Supplementary Fig. S5). For CO_2_ enrichment experiment, a total of 54,938 transcripts were differentially expressed between WL and CO_2_ enrichment. Based on adjusted *p*-value <10^-3^ and log2 fold change of two, 2,519 DEGs were identified, including 1,125 upregulated and 1,394 downregulated transcripts (Supplementary Fig. S5).

**Table 4:** Number of transcripts significantly differentially expressed in the four samples.

| Condition 1 | Condition 2 | Up-regulated | Down-regulated | Total | Category |
| --- | --- | --- | --- | --- | --- |
| BL | GL+RL+WL | 498 | 54 | 552 | BL-regulated |
| GL | BL+RL+WL | 320 | 1,656 | 1,976 | GL-regulated |
| RL | BL+GL+WL | 601 | 177 | 778 | RL-regulated |
| WL | BL+GL+RL | 127 | 2,534 | 2,661 | Either BL- or GL- or<br>RL-regulated |

To better understand the functions of DEGs, all the DEGs were mapped to the terms in KEGG database and compared with whole transcriptome background to search for genes involved in significantly pathways. Pathway enrichment analysis revealed that the annotated DEGs were mainly involved in ‘metabolic pathways’, ‘biosynthesis of secondary metabolites’, ‘oxidative phosphorylation’, ‘carbon metabolism’ and ‘porphyrin and chlorophyll metabolism’. The annotation shed light on the regulatory functions of light qualities on the specific processes, functions and pathways in *K. alvarezii*.

## Discussion

Intertidal seaweeds are subjected to rapidly changing environmental conditions. As a result, they have developed mechanisms that help them to cope with unfavourable disturbances or environmental stresses. These adaptation and acclimation responses contribute significantly to their survival and fitness, providing them with the plasticity necessary for a stationary life. This investigation highlights the understanding of gene expressions and responses of red alga *K. alvarezii* towards their light environment (BL, GL, RL and WL) and increasing CO_2_ level.

Photosynthesis begins with light absorbing, thus light harvesting is the first step in the photosynthetic process which mediated by pigment-binding proteins forming light-harvesting antenna systems. Light-harvesting complex (LHC) proteins constitute a large family of proteins, which includes chlorophyll *a/b*-binding proteins, fucoxanthin chlorophyll *a/c*-binding proteins, high light-induced proteins and early light-induced proteins (Green and Kuhlbrandt, 1995; Caron *et al*., 2001). Chlorophyll *a/b*-binding and fucoxanthin chlorophyll *a/c*-binding proteins were frequently reported to be transcriptionally repressed in response to light stress (Tonkyn *et al*., 1992; Maxwell *et al*., 1995; Teramoto *et al*., 2002). In this study, the amount of genes encoding light-harvesting proteins transcripts were significantly downregulated under different light qualities (BL, GL and RL) (Supplementary S5) and no significance difference between control and CO_2_ enrichment.

Fucoxanthin chlorophyll *a/c*-binding and early light-induced proteins were upregulated under GL. Meanwhile, similar responses of *K. alvarezii* were found under BL and RL that both fucoxanthin chlorophyll *a/c*-binding and high light-induced proteins were downregulated. Recent transcriptomic studies of stress responses highlighted fucoxanthin chlorophyll *a/c*-binding proteins were upregulated in responses to heat-, salt-, oxidative-, or light stress in both brown algae (Hwang *et al*., 2008; Dittami *et al*., 2009; Pearson *et al*., 2010). Similar observations were made concerning chlorophyll *a/b*-binding and fucoxanthin chlorophyll *a/c* ‐binding proteins in *Chlamydomonas reinhardtii* after high light treatment (Savard *et al*., 1996; Miura *et al*., 2004). It is reported that there is possible additional functions of these proteins. For example, fucoxanthin cholorophyll *a/c*-binding proteins were observed to be downregulated in a developmental mutant of the brown alga *Ectocarpus siliculosus* (Peters *et al*., 2008). The recently discovered “red lineage chlorophyll *a/b*-binding-like proteins” (RedCAPs) by Sturm *et al*. (2013) which was found to be restricted to the red algal lineage, reported to participate in the light (intensity‐ and quality‐) dependent structural remodelling of light-harvesting antennae in the red lineage. However, the existence and expression of RedCAPs in response to different light qualities and CO_2_ level in *K. alvarezii* remains unknown.

Chlorophyll and carotenoids in red algae is not that of a primary light absorber, this role is taken over by the phycobilisomes. Phycobilisome functions as primary light absorber in red algae and funnel the energy to the reaction center of PS II for conversion into chemical forms (Gantt *et al*., 2003). This complex contains two or three types of pigment-proteins known as biliproteins: phycoerythrin, phycocyanin and allophycocyanin. They differ in protein identity, chromophore type and attachment and their relative location in the phycobilisome complex (Blankenship, 2002). The phycobilisome absorbs light across the between 590-650 nm region of the solar spectrum that neither chlorophyll nor carotenoids are capable of absorbing light. Thus, organisms with phycobilisomes have greater accessibility to usable light within the visible spectrum and hence greater adaptability and light capturing capacity (Blankenship, 2002). Although RL had the significant impact on the growth rate of *K. alvarezii* as compared to those treated with BL and GL, however, the phycobilisomes showed no significant different expressions among the four light qualities. Different light spectral seemed to induce the same effect on phycobilisome contents in *K. alvarezii*. In contrast, many experiments (Figueroa and Niell 1990; Talarico and Cortese, 1993; Franklin *et al*., 2002; Godinez-Ortega *et al*., 2008) observed that the synthesis of chlorophyll, phycoerythrin, phycocyanin in the red algae have been shown to be influence by both irradiance and spectral composition. Such changes do not always imply statistical significance in differential expression analysis in transcriptome data as the differential analysis takes into account of overall depth of sequencing, read and fragment length, gene density in the genome, transcriptome splicing complexity and transcript abundance that could be vary for each sample (Trapnell *et al*., 2012). On the other hand, three DEGs tag to phycobilisome were identified between control and cultures treated with CO_2_ enrichment. Phycobilisome and phycoerythrin were upregulated while phycocyanin was downregulated under CO_2_ enrichment. The upregulation of photosystem proteins and phycobilisomes suggested that photosynthesis efficiency of *K. alvarezii* was enhanced by raising CO_2_ level.

The presence study has identified 38 gene tags of photosystems proteins (15 PS I and 23 PS II). Photosystems are functional and structural units of protein complexes involved in photosynthesis. The photosystem proteins showed no significant different expressions among the four light qualities. The most had the highest mRNA levels under RL (33) and the lowest under BL (18). It can be said that RL is more efficient than BL, GL and WL in term of photosynthetic efficiency in *K. alvarezii*. Meanwhile, out of 38 transcripts of photosystem proteins, 9 genes encoding of photosystem proteins were upregulated by raising CO_2_ concentrations, including 4 PS I proteins (ycf4, subunit III, reaction center subunit XI and chlorophyll A apoprotein) and 5 PS II proteins (D1, CP43 and CP47). These findings exhibited the importance of CO_2_ enhancing the quantum efficiency of photosystem I and II. Interaction between elevated CO_2_ levels and responses of seaweeds are still largely unknown. The contradictory results were obtained on growth and photosynthetic rates under CO_2_ enrichment (Gao *et al*., 1991, 1993; Garcia-Sanchez *et al*., 1994; Israel and Hophy, 2002; Zou, 2005). It is suggested that the functional PS II is increased under CO2 enrichment, therefore, in order to balance the stoichiometry of PS I and PS II, genes encoding subunits of PS I are simultaneously highly expressed to increase amount of PS I, improving the tolerance of plant to high CO2 levels.

Light-regulated gene expression mediated by photoreceptors has been study in plants, including algae (Ma *et al*., 2001; Jiao *et al*., 2005; see review Kianianmomeni and Hallman, 2014). We have identified two types of photoreceptors in *K. alvarezii*: phytochromes and cryptochromes. Phytochromes are red/far-red photoreceptors, measure the changes in light quality in the red and far-red regions of the visible spectrum, allowing plants to assess the quantity of photosynthetically active light (Franklin and Whitelam, 2005); cryptochromes are flavoproteins, function as blue light receptors that share sequence similarity to DNA photolyases, DNA enzymes that use blue light to repair UV-induced (Sancar, 2003). In the present study, phytochrome-like protein and cryptochrome transcripts were observed expressed under WL. The phytochrome gene is not expressed under BL, GL and RL. Thus we suggest that the expression of phytochrome genes in *K. alvarezii* seems to be triggered by others light qualities. In *Arabidopsis thaliana*, the expression of all photoreceptors genes has been demonstrated to be regulated by light or circadian clock (Reka, 2003). Similarly in brown alga *Saccharina japonica,* expression of photoreceptor genes seems to be triggered by all light qualities (BL, RL and WL) (Wang *et al*., 2013). In contrast, no phytochrome genes could be identified in the genomes of green alga *Volvox carteri* and *Chlamydomonas reinhardtii*, even though red‐ and far-red-regulated gene expression has been observed in these algae (Alizadeh and Cohen, 2010; Beel *et al*., 2012). The absence of phytochromes might indicate that other phytochrome photoreceptors such as animal-like cryptochromes, which absorb both blue and red light (Beel *et al*., 2012). On the other hand, the expression of cryptochrome genes was highly expressed in all light qualities and CO_2_ enrichment except under RL. Danon *et al*. (2006) and Lopez *et al*. (2011) reported that cryptochrome in plants are involved in the light-dependent gene expression, the light dependence mainly affect genes involved in the response to biotic/abiotic stress and regulation of photosynthesis. The presence of cryptochrome under GL is not surprising as cryptochromes are able to process GL, however, less effectively compared to BL (Folta and Maruhnich, 2007).

All the genes necessary to encode the enzyme involved in photosynthetic inorganic carbon fixation were identified (Supplementary Table S4). The data allow the identification of all enzymes necessary for the reductive pentose phosphate cycle, glyceraldehyde-3P => ribulose-5P (C_3_) and the C_4_-dicarboxylic acid cycle, phosphoenolpyruvate carboxykinase type (PCK) carbon fixation pathway, indicated the possibility of the existence of two photosynthetic pathways (C_3_ and C_4_) in *K. alvarezii*. Some experimental results suggest that the PCK-type is maximal in biomass production and CO_2_ fixation (Fravolini *et al*., 2002; Wang *et al*., 2012). Recent papers have reported the evidence for the operation of C_4_ photosynthesis as an alternative inorganic carbon-concentrating mechanism (CCM) in marine organisms (Tachibana *et al*., 2011). The expression levels of C_4_ enzymes, such as mdh, ast, ppc, pck and ppdk were higher than C_3_ enzymes, indicate C_4_ photosynthesis may function under atmospheric CO_2_ level. Most of the key enzymes of C_3_ such as glyceraldehyde 3-phosphate dehydrogenase (gapdh), pgk, and fructose-bisphosphate aldolase (fba) were expressed higher under CO₂ enrichment (Supplementary Table S6), however the difference is not significant. These results suggested CO₂ enhancement may alter carbon metabolism and lead to C_3_-type carbon metabolism in *K. alvarezii*. As CO_2_ level increase, the stomata plants increasingly close up and thus reduce the amount of water lost. In other words, plants significantly improve the water use efficiency as less water is transpired. Increase in the level of CO_2_, C_4_ plants show drastically reduced rates of photorespiration because CO_2_ is concentrated at the carboxylation site (RuBisCo) and is able to out compete molecular oxygen (Weber and von Caemmerer, 2010). The elimination of oxygenation reaction to a great extent and the loss of energy connected with the photorespiratory pathway is largely decreased (Heldt and Piechulla, 2011). For these reasons, the photosynthetic efficiency gap between C_3_ and C_4_ rapidly closes. As competing for the resources (water, nutrients and light) on the same patch, C_3_ is increasing outcompete the C_4_ metabolism. However, current understanding of the underlying mechanisms of the response of red algae to elevated CO_2_ concentrations is largely unknown; the demand of response of red algae to increasing CO_2_ warrants further research.

## Conclusion

To summarise, a total of 76,871 sequences were assembled using Trinity and 42,915 transcripts were assigned to functional annotation. 55.83% of transcripts were annotated through UniProt, provided important clues to the functions of many unknown sequences that could not be annotated using sequence comparison. Differential expression analysis revealed that only a small portion of genes (15.31%) were significantly expressed. The most DEGs were found under CO_2_ enrichment. Three different wavelengths were used: BL, GL and RL and compared to full spectrum (WL), similar gene expression patterns were found in term of photosynthesis aspect. The data presented suggesting light-regulated gene expression in *K. alvarezii* is not a single light response. On the other hand, CO_2_ enrichment leads to changes of photosynthetic pathway from C_4_ to C_3_ may contribute to the adaptation of this species to global CO_2_ changes. The alteration of photosynthetic pathways under CO_2_ enrichment highlights new questions to be addressed in subsequent work. This transcriptome study resulted in a number of new insights regarding the regulation of genes respond to different wavelengths of light and addition of CO_2_.

## Acknowledgements

The authors would like to thank the Ministry of Higher Education for granting a Fundamental Research Grant Scheme (FRGS) entitled “**Transcriptomic and carrageenan analyses of micropropagated *Kappaphycus* and *Eucheuma* seaweeds under different light regimes and carbon dioxide concentrations**”.

